# Identifying the oncogenic potential of gene fusions exploiting miRNAs

**DOI:** 10.1101/2021.10.20.465128

**Authors:** Venere S Barrese, Marilisa Montemurro, Marta Lovino, Elisa Ficarra

## Abstract

**Background:** It is estimated that oncogenic gene fusions cause about 20% of human cancer morbidity. Identifying potentially oncogenic gene fusions may improve affected patients’ diagnosis and treatment. Previous approaches to this issue included exploiting specific gene-related information, such as gene function and regulation. Here we propose a model that profits from the previous findings and includes the microRNAs in the oncogenic assessment. We present a classifier called ChimerDriver for the classification of gene fusions as oncogenic or not oncogenic. ChimerDriver is based on a specifically designed neural network and trained on genetic and post-transcriptional information to obtain a reliable classification.

**Results and discussion:** The designed neural network integrates information related to transcription factors, gene ontologies, microRNAs and other detailed information related to the functions of the genes involved in the fusion and the gene fusion structure. As a result, the performances on the test set reached 0.83 f1-score and 96% recall. The comparison with state-of-the-art tools returned comparable or higher results. Moreover, ChimerDriver performed well in a real-world case where 21 out of 24 validated gene fusion samples were detected by the gene fusion detection tool Starfusion.

**Conclusions:** ChimerDriver integrated transcriptional and post-transcriptional information in an ad-hoc designed neural network to effectively discriminate oncogenic gene fusions from passenger ones.

## Background

Gene fusions are one of the most common somatic mutations and are considered to be responsible for 20% of global human cancer morbidity [1, 2]. A gene fusion is a biological event where two independent genes fuse together to form a hybrid gene. In the most common case, one gene retains the promoter region and the other one provides the end of the hybrid gene. The former is referred to as 5p’ gene, while the latter is called 3p’ gene. The position where the break occurs is called breakpoint.

The advent of next-generation sequencing (NGS) and the development of fusion detection algorithms [3, 4, 5, 6] led to the discovery of hundreds of novel fusion sequences.

However, not all gene fusions are oncogenic. Indeed, some are genuinely expressed in normal human cells [7] or constitute passenger events [8]. At the same time, other gene fusions are considered to be responsible for a significant percentage of specific kind of tumors [9, 10, 11, 12].

A precise diagnosis of oncogenic gene fusions can inform therapeutics treatments [13, 14] and be used to predict prognosis, patient survival, and treatment response [2]. Additionally, focusing the research on a smaller number of putative oncogenic fusions a diagnosis could take less time; thus, the risks related to misdiagnosis and waiting may be significantly reduced for the patients.

However, discriminating between cancer-driver fusions and non-driver events is not a trivial task.

The first necessary step to solve this problem is performed by the fusion detection tools [3, 4, 5], that identify the candidate gene fusions relying on the sample’s reads, trying to reduce as much as possible the number of false positives (i.e., detected gene fusions that are not found in the sample in later lab validation). Additional studies proposed more sophisticated approaches based on machine learning (ML) techniques applied on top of fusion detection tools’ output. Specifically, Oncofuse [15] and Pegasus [16] are noteworthy and use protein domains of the fusion proteins to train the models and predict the oncogenic potential of a fusion. Undoubtedly protein domains are highly informative for the characterization of gene fusions. However, the use of such information as a feature for the ML model requires careful processing from scratch whenever the training database is updated with novel validated fusions.

Recently, previous works explored deep-learning (DL) techniques [17] and presented DEEPPrior [18], a DL model to perform gene fusion prioritization using amino acid sequences of the fusion proteins, based on a Convolutional Neural Network (CNN) and a bidirectional Long Short Term Memory (LSTM) network. Compared to the state-of-the-art tools, this approach is highly effective in accomplishing the classification task with the advantage of avoiding labor-intensive processing of the protein domains.

However, it is known that the oncogenic potential of a molecule depends not only on the sequence itself but also on the effect of post-transcriptional regulatory processes[19].

Transcription Factors (TFs) and micro-RNAs (miRNAs) play a decisive role in the transcriptional and post-transcriptional regulatory processes [20] and can contribute to determining the gene fusion outcome.

To date, most of the available tools exploit transcriptional information and common gene properties to accomplish this task, without considering the post-transcriptional regulators affecting the oncogenic processes.

Here, we present ChimerDriver, a new DL architecture based on a Multi-Layer Perceptron (MLP) which integrates gene-related information with miRNAs and TFs including then in the model transcriptional and post-transcriptional regulative information. Indeed, ChimerDriver exploits the knowledge about TFs and miRNAs targetting each of the genes involved in the fusion to perform gene fusion classification.

ChimerDriver was tested on multiple publicly available datasets and exhibited better classification performance with respect to the state-of-the-art tools. In the end, post-transcriptional regulators confirm the central role in the discovery of oncogenic processes and miRNAs, in particular, they are a precious source of information to improve the prediction of the oncogenic potential of gene fusions.

In the following, the proposed method is illustrated alongside with the results in Results section. The discussion and conclusion are reported in Discussion and Conclusions sections, respectively, while a detailed description of model, its architecture and the input datasets is provided into the Methods section.

## Results

This section presents an overview of ChimerDriver and of the datasets used in the training, testing, and validation phases. Additionally, the results obtained with ChimerDriver and the comparison with the state-of-the-art tools are discussed. In the end, a case study in which ChimerDriver was applied on a pair of well-known datasets is presented.

A detailed description of ChimerDriver architecture and of the used datasets is provided in the Methods section.

### Architecture overview and results on the test set

ChimerDriver is based on a feed-forward multi-layer perceptron (MLP). The input feature set combines structural properties of 5p’ and 3p’ genes with a comprehensive profiling of transcriptional and post-transcriptional regulators (TFs and miRNAs).

In details, we considered as input features the retained percentage of genes, the strand and the relevance in cancer of both 5p’ and 3p’ genes. Additionally, the oncogenic role of each gene was taken into account. This information was extracted by Cancermine [21], a database which classifies genes as drivers, oncogenes or tumor suppressors. The label ’other’ is used when none of the above options apply.This feature contributes to the assessment of the functional profiling of the gene fusion. TFs and Gene Ontologies (GOs) were also included in the feature set due to their importance in assessing the oncogenic potential of gene fusions[15]. Finally, we added miRNAs to consider post-transcriptional regulation, which influences the translation of gene fusions and therefore their actual oncogenic activity. Specifically, we considered all miRNAs which target 5p’ and 3p’ genes according to Targetscan database[22].

Due to the high complexity of the model and extensive amount of features, a feature selection process has been performed on the input feature set. The model was trained on 1765 gene fusions, obtained from COSMIC, Catalog of Somatic Mutations in Cancer [23] and from Babicenau et al. work [24]. Given each gene fusion’s breakpoint, the aforementioned features are extracted and then fed to the MLP. For more details about the feature selection and the data used, refer to the Methods section.

The model was cross-validated on 10 folds using different learning rates and dropout values to find the classification task’s best parameters. The model reached an average f1-score of 0.98 on our training set with different combinations of learning rate and dropout values.

According to the cross-validation results, the best network configuration was characterized by four layers with respectively 512, 256, 128, 64 nodes. For each node, the associated activation function was the tanh. The best learning rate was found to be 0.01, while the best dropout value applied to each layer was 0.2. Therefore the model was tested on 4877 gene fusions. 2623 oncogenic gene fusions were retrieved from the work of Gao et al.[25] and the remaining 2254 were gene fusions found in healthy tissues and reported by Babicenau et al. [24].We ensured that the test samples are entirely independent from the training samples. The model returned a 0.83 f1-score and 96% recall when tested on this set of gene fusions.

### miRNA impact on the classification performance

The miRNA features were extracted from TargetScan [22], a popular database that maps gene-miRNA pairs providing various kinds of information. We mainly focused on the probability that the miRNA would target the specific gene during post-transcriptional regulation. This value was extracted for both 5p’ and 3p’ genes and it is intended to represent the involvement of the miRNAs in the gene fusion processes. In figure 1 we highlight the impact of the miRNA features in the classification by displaying the confusion matrices including and excluding miRNAs from the evaluation. The impact of miRNAs is particularly evident when looking at the number of false-negative gene fusions, which is almost doubled when miRNAs are not taken into consideration. By including miRNAs in the classification task the recall value increases from 93% to 96%.

**Figure 1:**
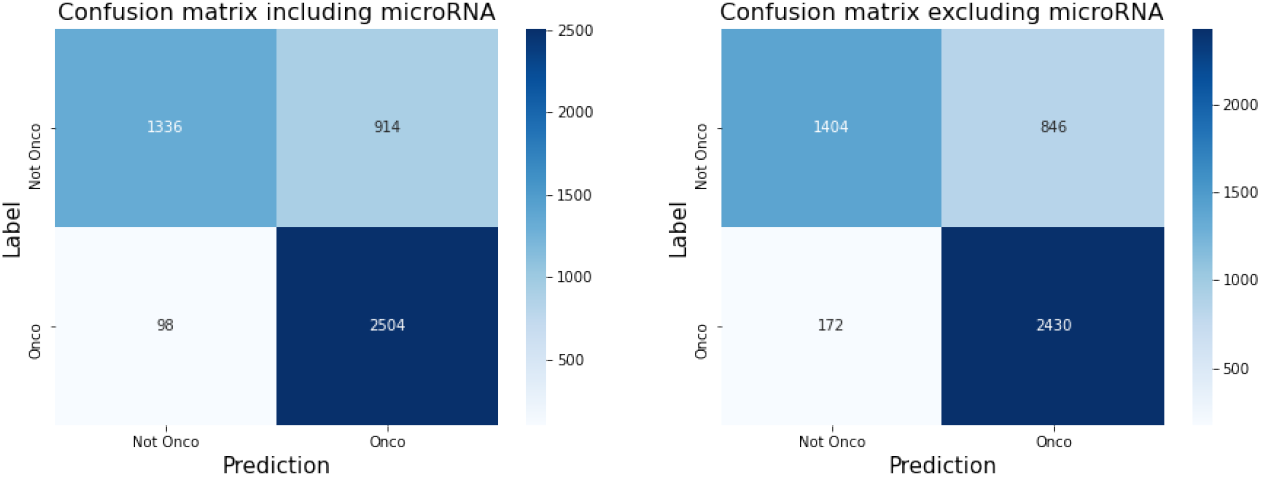
Confusion matrices reporting the MLP results including miRNAs (on the left) and excluding miRNA features (on the right).

### Comparison with state of the art

ChimerDriver performances were compared to the ones reported by three related works: Oncofuse[15], DEEPrior[18], and Pegasus[16]. To compare the results in the most unbiased way, the experimental conditions of the three tools were reproduced and ChimerDriver was applied.

#### Oncofuse

To test the robustness of the proposed method, we extrapolated the training set and testing set used by Oncofuse [15]. Those samples were used to train and test our model and then to compare Oncofuse and ChimerDriver performances.

Oncofuse training samples were extracted from TICDB [26], a curated database that contains gene fusions found in tumor samples, and from a collection of fusion genes [27], and read-through transcripts [28] found in normal cells named NORM-RTH. Oncofuse’s authors then built the oncogenic testing set by merging oncogenic gene fusions from CHIMERDB [29] and NGS, respectively oncogenic fusions predicted by gene fusion detection tools and fusions discovered and published in NGS studies about cancer [30, 31, 32, 33]. On the other hand, not oncogenic testing samples were taken from Refseq [34] and CGC [35], two databases that report unbroken gene fusions. In particular, the samples that belong to CGC involve unbroken oncogenic genes.

All the previously listed features (see Methods for details) were processed and gathered, except for the two features related to the retained percentage of genes. These features could not be considered in the evaluation since the provided dataset omitted the breakpoint information.

ChimerDriver model was tailored to this comparison. We obtained 281 input features: the strands and the involvement in oncogenic processes of both 5p’ and 3p’ genes, 93 TFs, 155 miRNAs, and 30 GOs. The maximum number of epochs was set to 50, and the number of nodes per layer was 256, 128, 64, and 32 (the associated activation functions were the relu, sigmoid, relu, and sigmoid respectively). The learning rate was fixed to 0.03, while the dropout value applied to each layer was 0.4.

Figure 2 shows the comparison of the classification results obtained by ChimerDriver and Oncofuse. Precisely, the green bars correspond to the results reported by Shugay M. et al. [15] for Oncofuse performances. In blue, the results obtained by ChimerDriver are displayed.

**Figure 2:**
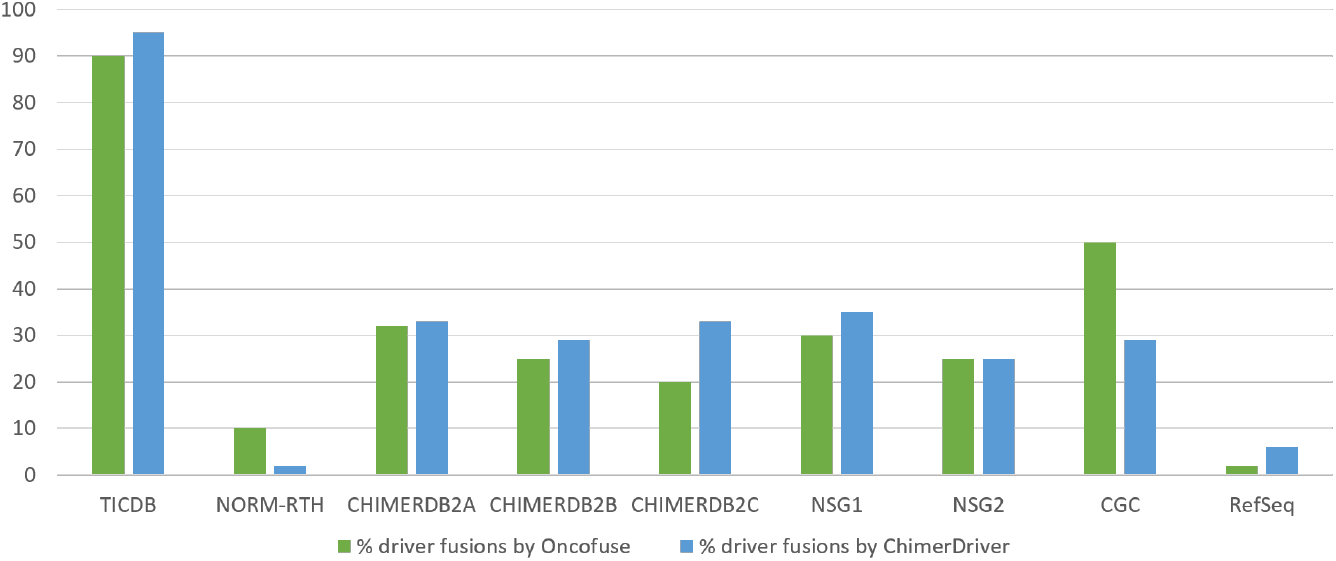
The green bars correspond to the results reported by Shugay M. et al. in their paper. In blue the results obtained by ChimerDriver are displayed.

The results of ChimerDriver, when trained and tested on the samples provided by Oncofuse, were able to outperform the ones illustrated in the original paper. Similarly to the research conducted by Shugay et al., the results for each database are displayed separately. The bar diagram shows the percentage of driver gene fusions detected by the model.

Indeed, ChimerDriver results reported in 2 show high performances for the training set, outperforming the ones obtained by Oncofuse. 95% of TICDB samples were correctly classified as driver gene fusions by ChimerDriver as opposed to the assumed 90% reported by Oncofuse, furthermore 2% of the NORM-RTH samples were incorrectly classified as driver gene fusions by ChimerDriver as opposed to the assumed 10% reported by Oncofuse.

ChimerDriver successfully outperformed Oncofuse in the oncogenic gene fusion databases used as a test set, namely ChimerDB2A, ChimerDB2B, ChimerDB2C, NGS1 and NGS2. ChimerDriver identified more or a comparable amount of oncogenic gene fusions in each database with respect to Oncofuse, correctly classifying about 1/3 of the samples.

ChimerDriver minimized the number of detected driver fusions of unbroken oncogenic genes, identifying a lower number of driver gene fusions in CGC database, as additional test set.

When tested on the not-oncogenic samples in RefSeq database, Chimerdriver returned a slightly higher number of driver gene fusions.

In general, we may conclude that even without the information on the retained percentage of genes, ChimerDriver outperformed Oncofuse in the great majority of cases.

#### DEEPrior

DEEPrior is a DL-based classifier which performs gene prioritization using protein sequences obtained from the gene fusion samples. Its architecture is based on a CNN and an LSTM network. It was trained on a dataset extracted from COSMIC [23], and Babicenau et al.’s study [24] and tested on part of the oncogenic gene fusion collection validated by Gao et al. [25]. DEEPrior reconstructs the protein sequences from gene fusion breakpoint information and assigns to each gene fusions an oncogenic score defining its oncogenic probability. Gene fusions are ordered according to the oncogenic score and highly scored fusions are prioritized as drivers. In this sense, DEEPrior main aim consists in providing a reliable classification prediction (oncogenic or not) according to the oncogenic score.

We trained and tested ChimerDriver on DEEPrior training set and test set (*Dataset 2* in DEEPrior paper). As a result, ChimerDriver correctly classified 96% of oncogenic gene fusions from the test set. On the contrary, DEEPrior prioritized as driver the 32.48% of gene fusions found in the test set. Since DEEPrior aims at classifying only highly probable oncogenic fusions, the percentage of prioritized gene fusions is not directly comparable with the classification performances obtained with ChimerDriver. ChimerDriver provides a classification result for each gene fusion, while DEEPrior classifies a very small percentage of gene fusions in the dataset.

We can conclude that ChimerDriver approach exploits different sources of information (TFs, GOs, miRNAs) while DEEPrior focuses on identifying the oncogenic potential of a gene fusion through its protein sequence without considering the effect of post-transcriptional regulators.

At the same time, ChimerDriver ensures a less computationally intensive approach in the training phase compared to DEEPrior.

#### Pegasus

To further assess ChimerDriver classification performances, we took into account Pegasus [16], a state-of-the-art tool for gene fusion detection and classification purposes. Pegasus exploits a traditional machine learning model for the prediction of driver gene fusion, namely a gradient tree boosting algorithm.

Also in this case, ChimerDriver was trained and tested on the gene fusion samples used to develop and validate Pegasus.

We observed that the training dataset was strongly unbalanced towards the negative samples, comprising of over 9923 negative samples out of 10162 gene fusions. In order not to penalize the MLP architecture which is particularly sensible to class unbalance, we lowered the number of negative gene fusions to 239, namely the number of positive samples.

ChimerDriver was cross-validated on 10 folds using the aforementioned training samples. It should be noted that, as a result of balancing the classes, the model was given a fairly small number of training examples. In the end, the f1-score was equal to 0.89 with a learning rate and dropout respectively equal to 0.001 and 0.

Pegasus’s test set accounted for 78 gene fusions, 39 oncogenic and 39 not oncogenic respectively. According to Pegasus authors, the curated subset of 39 oncogenic gene fusions were almost entirely correctly classified by Pegasus that reported 0.97 of AUC and 0.95 of AUC for the not oncogenic samples.

Pegasus intently selected as negative examples 39 not oncogenic gene fusions containing at least a tumor suppressor or an oncogene. The rationale is that these gene fusions would be most challenging for a classification task. ChimerDriver correctly classified 27 out of the 39 not oncogenic gene fusions enforcing the notion that the model is able to generalize even on not oncogenic gene fusions. On the other hand, the oncogenic test samples represented a more difficult classification task for ChimerDriver, which detected 17 oncogenic gene fusions. It should be noted that ChimerDriver model was originally trained and tested on a wide variety of gene fusions proving its ability to learn and generalize well when given a fair amount of examples. On the contrary, since Pegasus was developed and refined on particular tissues, a reduced number of samples is used as a training set.

In our opinion, the small number of samples in the Pegasus training set negatively impacted the ChimerDriver training phase, which benefits from a wider number of gene fusions. Therefore, ChimerDriver performances, when trained and tested on Pegasus datasets, are negatively affected, hindering the likelihood of reaching the outcome reported by the Pegasus authors.

### Case study

Finally, to assess ChimerDriver’s performances in a clinical context, we selected two well-known studies: 6 breast cancer samples [36] and 4 prostate cancer samples [37] in which 24 gene fusions are reported to be experimentally validated. The samples are all RNA-seq data. We processed them with STAR-fusion [38] to identify which gene fusions were found in these samples by a standard and accurate fusion detection tool. 21 out of the 24 validated gene fusions were actually detected with STAR-fusion and subsequently processed with ChimerDriver to confirm the ability in correctly detecting oncogenic gene fusions in a real-world case. Figure 3 shows the results of this assessment. Specifically, the gene fusions marked in gray were not detected by STAR-fusion hence were not available to ChimerDriver for further processing. The training dataset and the training parameters are described in detail in the Methods section like the ones generally used in the ChimerDriver training procedure. On the 21 samples, ChimerDriver wrongly classified as not oncogenic the three oncogenic gene fusions marked in orange. By inspecting the oncogenic role of 5p’ and 3p’ genes, and also the retained percentage in the gene fusion, a possible explanation for the wrong classification could be hypothesized. Concerning the ACACA-STAC2 gene fusion, no information on the involvement of any of the two genes was provided to the algorithm. So, although most of the portion of both genes was retained after the gene fusion event, ChimerDriver was probably unsure about their role in oncogenic processes. As for the GLB1-CMTM7 fusion, the algorithm was aware that the latter gene is involved in tumor suppression, on the other hand the retained percentage of CMTM7 was less than 45%. This probably led to the conclusion that there was not enough gene left in the gene fusion to cause issues. Similarly, in the CPNE1-PI3 fusion the percentage of retained genes (respectively 25% and 40%) was probably too low to label the gene fusion as oncogenic even if the genes were associated to the roles oncogenic and driver respectively. Finally, ChimerDriver correctly classified the 18 remaining gene fusions as oncogenic. Hence, ChimerDriver correctly classified 18 out of 21 oncogenic gene fusions, demonstrating that the specifically designed neural network is proficient in learning and generalizing from a consistent number of gene fusion samples. Moreover, the information gathered from the different sources and provided to the tool as features proved to be particularly effective in discerning between oncogenic and not-oncogenic fusions even in a realistic circumnstance.

**Figure 3:**
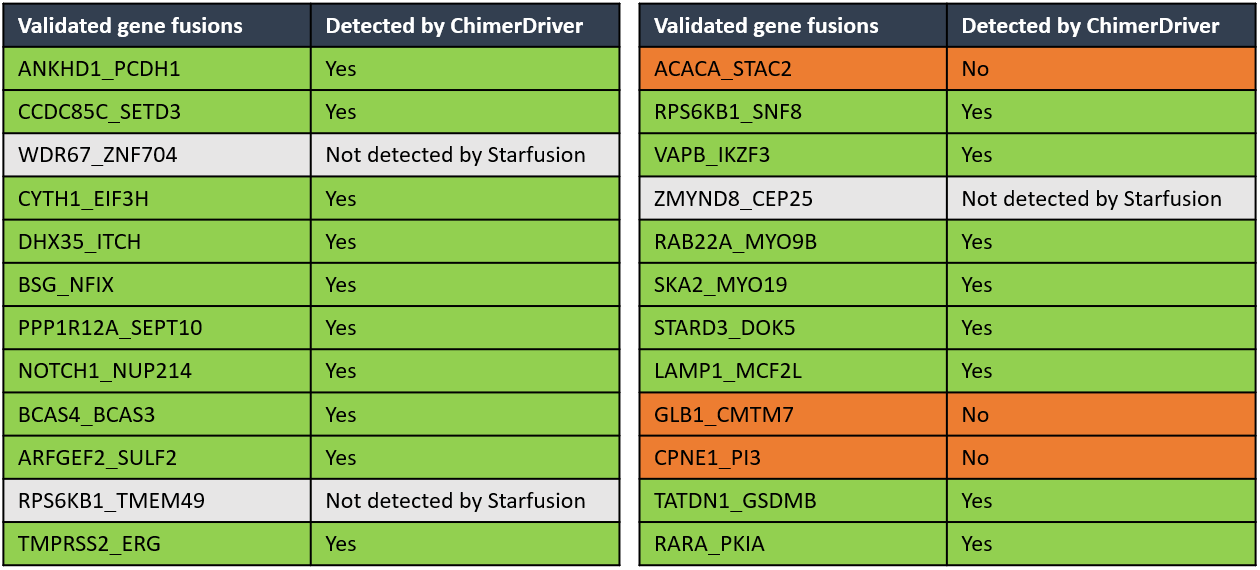
The 24 oncogenic gene fusions validated in prostate and breast tumor samples are reported. STAR-fusion did not detect the three gene fusions marked in gray hence were not available to ChimerDriver for further processing. ChimerDriver correctly classified as oncogenic 18 out of the 21 available gene fusions.

## Discussion

Identifying oncogenic gene fusions is of crucial importance in cancer detection and prognosis. To date, state-of-the-art tools exploit transcriptional and GOs information, without considering the post-transcriptional regulators in predicting the oncogenic potential of a gene fusion. Here, we presented ChimerDriver, a novel tool to accomplish the aforementioned task exploiting transcriptional and post-transcriptional regulators. In details, ChimerDriver focuses on miRNAs post-transcriptional effect as a key feature to performe the prediction.

ChimerDriver is based on an ad-hoc designed neural network embedding miRNAs, transcription factors, gene ontologies, and gene-specific information to predict gene fusions’ oncogenic potential. The model is stable and exhibits excellent classification performance (f1-score = 0.98).

We tested our classifier against three state-of-the-art tools: Oncofuse, DEEPrior, and Pegasus.

With respect to Oncofuse, we introduced post-transcriprional regulation to perform the classification and, as a result, ChimerDriver outperformed Oncofuse in the great majority of tested cases.

In particular, ChimerDriver performed better than Oncofuse on the test set, correctly classifying as oncogenic about 1/3 of the oncogenic gene fusions. ChimerDriver identified a comparable or higher amount of oncogenic gene fusions outperforming Oncofuse results in each of the positive test cases. ChimerDriver minimized the number of detected driver fusions in ’unbroken oncogenic genes’ (negative testing samples) extracted from CGC compared to Oncofuse. This result confirmed the ability of ChimerDriver in generalizing and taking advantage of the given set of features to make a correct prediction. As previously presented in the Results section about Pegasus comparison, this statement is true even when the samples contain an oncogene or a tumor suppressor. ChimerDriver returned a slightly higher number of oncogenic gene fusions than Oncofuse when tested on RefSeq database of ’unbroken not-oncogenic genes’. We recall that the breakpoint information was not available in Oncofuse datasets. Therefore, to perform an unbiased comparison with Oncofuse, the breakpoint information was neglected by ChimerDriver model. Consequently, the percentage of driver gene fusions detected by ChimerDriver on RefSeq was slightly higher than expected probably because the tool could no profit from the breakpoint information.

ChimerDriver also outperformed DEEPrior in terms of the number of classified gene fusion. In particular, ChimerDriver correctly identified 96% of oncogenic gene fusions in the dataset used to test DEEPrior, which prioritized as oncogenic only 32.48% of the samples. It should be noted that the goals of DEEPrior and ChimerDriver are slightly different. The first performs a prioritization of gene fusions, returning those with an oncogenic probability greater than a threshold (typically 80%). ChimerDriver instead performs an immediate classification of each gene fusion by integrating transcriptional and post-transcriptional features in the assessement. The final outcome of ChimerDriver is remarkable in terms of number of oncogenic samples that were correctly classified while also enlightening because it stresses the extent in which miRNAs are involved in the oncogenic processes of gene fusions.

Moreover, the performances of ChimerDriver were compared to the ones reported by Pegasus authors. According to their research, the latter was able to correctly classify almost all of the test samples. After training and testing ChimerDriver on the gene fusions provided by the authors, it was observed that the number of detected oncogenic samples was lower than the results reported by Pegasus. As already stated in the Results section, the number of training samples was lowered in order to balance the oncogenic and not oncogenic classes. However, the limited number of samples processed by ChimerDriver in the training phase has probably inhibited the neural network from learning efficiently. In addition, Pegasus’s authors specify that the negative validation samples included at least one oncogene or tumor suppressor. We remind that, to make a prediction, ChimerDriver also relies on the role of each gene in oncogenic processes (e.g., driver, oncogene, or tumor suppressor), making the classification task particularly arduous to tackle. In addition, Pegasus and consequently ChimerDriver were trained on a reduced number of samples, thus impacting ChimerDriver performances. Nevertheless, ChimerDriver correctly classified most of the not oncogenic gene fusions enforcing the notion that the model is capable of generalizing well in this situation.

In this work, we focused on the integration of information coming from different databases to improve the current state-of-the-art research on classifying oncogenic gene fusions. Additionally, a neural network was specifically designed for this task. However, the main contribution of the present work is the introduction of miRNAs in the classification model. In fact, despite miRNAs role in determining the oncogenic potential of gene fusions has been demonstrated, they had never been considerated in such a task. In the present work, we showed that they could significantly improve the model performance. In particular, they halved the number of false negatives and improved the recall of the model. We can conclude that miRNAs, being involved in the regulation of gene fusion-related protein, are a promising indicator of the oncogenic potential of gene fusions.

The main limitation of the proposed method is that some gene fusions are mis-classified. To better investigate ChimerDriver classification with respect to the Cancermine [21] role, we reported in Figure 4 the distribution of the Cancermine roles (e.g. tumor suppressor, driver, oncogene, other) for 5p’ gene (Figure 4a) and 3p’ gene (Figure 4b). In addition, test set samples are divided in each role according to the classification results (false positives (FP), false negatives (FN), true positives (TP) and true negatives (TN)). TP samples are characterized both for 5p’ ad 3p’ genes by a prevalence of suppressors and oncogenes. On the contrary, TN mostly refer to the ’other’ CancerMine role. As a consequence, FP samples could consist of oncogenes (in particular for 3p’ gene) and FN samples are hardly ever related to tumor suppressors, drivers, or oncogenes. In this sense, FP and FN samples reflects ChimerDriver behaviour on TP and TN respectively. In a clinical context, FN misclassified samples are unlikely to be tested for in lab validation, since most of them involve genes with not a specific oncogene/tumor suppressor role. FP samples instead would have been considered for an experimental validation, that in the the end would exclude them from oncogenic fusions. However, laboratories would still benefit from a selection of putative oncogenic gene fusions.

**Figure 4:**
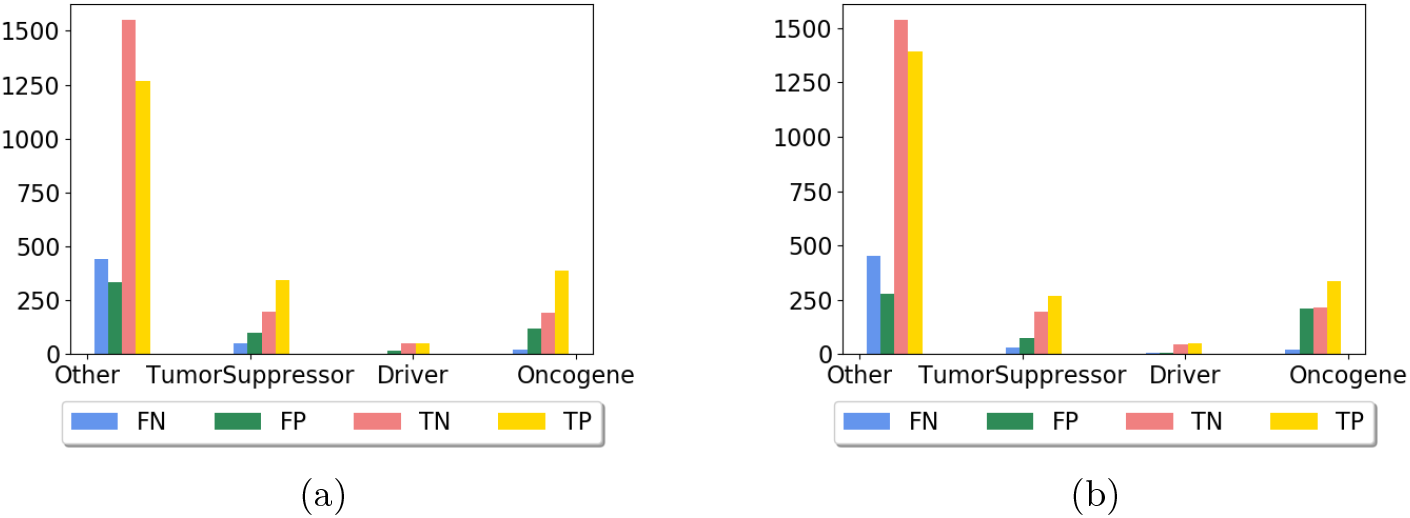
Here we report the distribution of the false positives (FP), false negatives (FN), true positives (TP) and true negatives (TN) regarding Cancermine information for both 5p’ and 3p’ genes(respectively Figure 4a) and 4b)). Noticeably, FPs are never tumor suppressors, drivers or oncogenes.

## Conclusions

Gene fusions are a common mutation that is nowadays known to be responsible for about 1/5 of human cancers. It is of the uttermost importance to correctly identify gene fusions to improve cancer detection and prognosis. Considering that the state-of-the-art tools exploit transcriptional and gene information neglecting post-transcriptional regulations, we combined this knowledge and established the value of miRNAs in achieving superior classification performances.

To conclude, we presented ChimerDriver, a novel and stable DL architecture based on a Multi-Layer Perceptron (MLP) which, for the first time, combines gene-level features with TFs and miRNAs targetting the gene fusion to perform its classification and prioritization.

ChimerDriver was trained and tested on a consistent number of gene fusions. The final results highlight the impact of miRNAs in the evaluation of the oncogenic potential of gene fusions. We can infer that the inclusion of miRNAs represents a valuable advantage in the identification of oncogenic gene fusions.

ChimerDriver can become a valuable tool for research laboratories to predict the oncogenic potential of gene fusions. Indeed, the expensive validations could be targeted cost-effectively with this easy-to-use tool; additionally, it may speed up identifying novel and potentially oncogenic gene fusions, allowing for better diagnosis, classification, and treatment of cancer patients.

## Methods

The oncogenic potential of gene fusions was assessed through an MLP to approach this challenging task. The MLP was believed to be a suitable method since a simple yet effective configuration can characterize it. The process of adjusting the hyper-parameters of the MLP allowed for fruitful research for the best model to achieve the highest possible performances.

The chosen dataset used for training and testing the model is broad and extensively described in Dataset subsection.

The features used by the MLP were carefully constructed to obtain the highest possible degree of information concerning the gene fusion samples. Subsequently, the high number of features was reduced through the Random Forest feature selection technique.

### Feature selection

All the available features come from multiple sources, and they are related to different characteristics of the gene fusions.

The first five features are obtained from gene fusion structure, and Cancermine [21], a literature-mined database of drivers, oncogenes, and tumor suppressors in cancer. Two features correspond to the retained percentage of 5p’ and 3p’ genes in the gene fusion, given the breakpoint coordinates. One additional feature controls for the strands of 5p’ and 3p’ genes, and it is equal to 1 if the two strands are concordant (the two genes transcribe in the same direction), 0 otherwise. The remaining two features correspond to the nature of each gene according to Cancermine [21]: ’Oncogenic’, ’Driver’, ’Tumor suppressor’ or ’Other’ when none of the above options apply.

Other studies [15] have already covered the impact of TF [39] and GOs [40] in the gene fusion classification, which proved to be extremely useful for this purpose. Therefore, TF and GOs have been included in the ChimerDriver model. Specifically, a set of 181 TFs was extracted from the ENCODE database [39] and only those related to the gene in the 5p’ position were considered.

Additionally, since each gene can be involved in many different GOs, all of them have been selected. This approach resulted in an extensive amount of GOs to consider, that is, 5125 features.

Besides, our main contribution consists of including miRNAs post-transcriptional regulation in the model. Specifically, all miRNAs predicted to target all 5p’ and 3p’ genes have been considered. This information was extracted from TargetScan, a popular state-of-the-art database that predicts biological targets of miRNAs by searching for the presence of sites that match the seed region of each miRNA[22], reporting for each miRNA all possible target genes. A set of 333 miRNAs was obtained by investigating the probability of both genes belonging to the gene fusion. In case of ambiguity, only the highest probability was retained.

The final feature set was considerably lengthy. Thus, we performed feature selection to reduce the 5644 total features to a more reasonable number. The chosen feature selection method was the random forest in which the number of features was lowered according to a threshold. The higher the threshold, the lower the number of retained features. For this study, the already stated threshold was kept in the range 0.0001-0.0005.

### Dataset

We retrieved 1765 samples for the training set, 1059 labeled as oncogenic gene fusions and the remaining 706 as not oncogenic. On the other hand, the testing set consisted of 2623 positive samples and 2254 negative samples. A set of 156 gene fusions (122 positives and 34 negative samples) was used in combination with 200 randomly selected samples of the training set as a validation set during the neural network training phase. The processing performed by ChimerDriver to build the features for these samples caused a modest amount of gene fusions to be discarded. It was due to unrealistic values obtained from the calculation of the percentage of the retained gene due to occasional errors in retrieving the correct breakpoint value or strand of a limited amount of genes. However, the majority of the samples adopted in this study are in common with DEEPrior work[18].

#### Training set

The oncogenic samples of the training set were extracted from COSMIC (Catalog of Somatic Mutations in Cancer). This popular database includes information on gene fusions involved in solid tumors and leukaemias. [23] The chosen 1059 oncogenic gene fusions were already experimentally validated. Moreover, the exact breakpoint positions were provided for each of them. On the other hand, the 706 not oncogenic gene fusions were reported by Babicenau et al. [24] and detected by a gene fusion detection tool in non-neoplastic tissues.

#### Test set

To build the test set, we used the database provided by Gao et al. [25] which is the result of three fusion detection tools applied on the TCGA database. Among these samples, the authors kindly provided validated gene fusions upon request. These samples were the ones for which WGS data were available. From this collection, we extracted 2622 oncogenic gene fusions. Besides, we incorporated a comparable number of negative samples to better attest our tool’s performances. The set of 2254 not oncogenic gene fusions was reported by Babicenau et al. [24]. These gene fusions were found in healthy tissues.

### Model architecture

As previously stated, an MLP was explicitly designed to evaluate the oncogenic potential of the gene fusions. Four layers characterized the final model. For each layer, the number of nodes was respectively: 512, 256, 128, 64.

The activation functions were varied to check which configuration would return the highest performances. Different combinations of the following functions were used: Sigmoid, Tanh and Relu.

Other parameters that have been varied were the learning rate, the dropout, and the number of epochs.

- Learning rate: 0.0001, 0.001, 0.01
- Dropout: 0, 0.1, 0.2, 0.3, 0.4
- Number of epochs: 500 - 1000

The final tool can either take advantage of an early stopping module that stops the training when the accuracy does not improve for 50 consequent epochs or train for a fixed number of epochs.

As already stated, the tool’s validation set is a combination of 156 gene fusions and 200 randomly selected samples from to the training set. The goal was to validate the model on a set that included information coming from a different source concerning the training set. In this case, the 200 samples used for validation were not considered in the training phase.

## Abbreviations

TF: transcription factors
GO: gene ontologies
miRNAs: microRNAs
MLP: multilayer perceptron

## Ethics approval and consent to participate

Not applicable.

## Consent for publication

Not applicable.

## Availability of data and materials

Source code, datasets and a minimal documentation are available on GitHub: https://github.com/veneresabrina/ChimerDriver.

## Competing interests

The authors declare that they have no competing interests.

## Funding

Not applicable.

## Competing interests

The authors declare that they have no competing interests.

## Author’s contributions

Conceptualization, V.S.B., M.M., M.L. and E.F.; methodology, V.S.B., M.M., M.L. and E.F.; software, V.S.B., M.M.; validation, V.S.B., M.M.; formal analysis, V.S.B., M.M., M.L. and E.F.; data curation, M.L. and V.S.B.; writing—original draft preparation, V.S.B., M.M.; supervision, M.L. and E.F.; funding acquisition, E.F.

## Acknowledgements

Computational resources were provided by HPC@POLITO, a project of Academic Computing within the Department of Control and Computer Engineering at the Politecnico di Torino (http://www.hpc.polito.it).

## Notes

### Competing Interest Statement

The authors have declared no competing interest.

